# Leptospermum extract (QV0) suppresses pleural mesothelioma tumour growth *in vitro* and *in vivo* by mitochondrial dysfunction associated apoptosis

**DOI:** 10.1101/2022.12.06.519377

**Authors:** Huaikai Shi, Le Zhang, Ta-Kun Yu, Ling Zheng, Helen Ke, Ben Johnson, Emma Rath, Kenneth Lee, Sonja Klebe, Steven Kao, Karl Lijun Qin, Hong Ngoc Thuy Pham, Quan Vuong, Yuen Yee Cheng

## Abstract

Pleural mesothelioma (PM) is a highly aggressive, fast-growing asbestos-induced cancer with limited effective treatments. There has been an interest in using naturally occurring anticancer agents derived from plant materials for the treatment of PM. However, it is unclear if aqueous extract from the *Leptospermum polygalifolium* (QV0) has activity against PM. Here we investigated the anti-cancer property of QV0 *in vitro* and *in vivo*.

Animals treated with Defender^®^ (QV0 dietary supply) exhibited a reduced tumour size over 30 days, which was associated with an average extended of seven days mouse life. There was no liver toxicity, nor increased blood glucose post-treatment in animals treated with Defender®. Moreover, QV0 suppressed the growth of 13 cancer cell lines in a dose-dependent manner, effective at concentrations as low as 0.02% w/v. This response was found to be associated with inhibited cell migration, proliferation, and colony formation, but without evident cell cycle alteration. We observed mitochondrial dysfunction post QV0 treatment, as evidenced by significantly decreased basal and maximal oxygen consumption rates. Significantly enhanced tumour apoptosis was observed in the Defender®-treated animals, correlating with mitochondrial dysfunction. To the best of our knowledge, this study constitutes the first demonstration of an improved host survival (without adverse effects) response in a QV0-treated PM mouse model, associated with an evident inhibition of PM cell growth and mitochondrial dysfunction-related enhancement of tumour apoptosis.

**Importance:** A major problem with cancer chemotherapy or immunotherapy is the severe adverse effects associated with normal tissue damage. PM is known to be treatment resistant and has poor a prognosis, therefore new therapeutic treatment options are urgently needed. In the present study, we explored the potential utility of a *Leptospermum* extract (QV0) as a treatment option for mesothelioma. We demonstrated for the first time that QV0 exhibits an anti-tumour response in mesothelioma, without any associated adverse effects observed in the PM mouse model. These findings provide a rationale for early-stage clinical trials. We anticipate that prospective translational research will lead to the clinical implementation of a novel QV0-based treatment strategy that will ultimately benefit PM patients.

## Introduction

Pleural mesothelioma (PM) is an aggressive thoracic malignancy with a poor prognosis and high symptom burden that is caused by previous exposure to asbestos. The current treatment options for PM patients include cisplatin and pemetrexed^1^, with the possible addition of bevacizumab to chemotherapy^2^, or combination immunotherapy with ipilimumab and nivolumab^3,4^. Despite the recent advancement in PM treatment with the introduction of immunotherapy, the average survival time of PM patients remains poor, with a median survival of around 18 months^5^. Novel treatment agents and approaches are desperately needed to improve PM patient survival outcomes.

Natural products including plants, microbial products, and marine sources have provided key substrates in the production and development of anti-cancer drugs for decades^6^. There remains great interest in the potential for natural products to produce new anti-cancer therapies. Unlike the significant toxic side effects associated with the standard chemotherapy- and immunotherapy-based cancer treatment options, novel drug candidates developed from natural products induce minimal toxicity on healthy non-malignant cells^7^. A variety of natural substances have been tested *in vitro* and in PM animal models. These include various polyphenolic compounds^8^ such as curcumin^9^, resveratrol^10^ and quercetin^11^, as well as extracts from plants such as the artichoke leaf (Cynara scolymus)^12^, olive leaf (Olea europaea L.)^13^, Glychyrrhiza inflata^14^, Filipendula vulgaris^15^ and microbial products such as Maunomycin A^16^ and JBIR-23^17^.

The present study represents the first report on the anticancer activity of a specific extract from the manuka honey tree, *Leptospermum polygalifolium* (QV0, P116949.AU), in PM. In animal experiments, QV0 delivered as a dietary supplement using manuka honey as a base (Defender®) also demonstrated tumour suppressive activity without clinical, biochemical, or anatomical evidence of toxicity. These findings provide a rationale for prospective translational research aimed to facilitate the clinical implementation of a QV0-based PM treatment.

## Results

To investigate whether QV0 has anti-cancer property, we first treated the xenografted mesothelioma (MSTO) mice with a daily oral administration of Defender®, a dietary supply of QV0 at 0.25mg/g/day for 30 days (Figure 1A). Tumour growth was indicated as cell counts and were monitored using IVIS imaging system following 9, 16, 23 and 30 days post-tumour implantation, respectively (Figure 1B). We found that the tumour volume increased progressively in control animals, with approx. 7.67e07 tumour cells measured at day 16 after tumour implantation. In comparison, mice treated with Defender® exhibited a significant reduction in tumour volume, with approx. 3.16e07 cells measured at day 16; representing a 41 % inhibition of tumour growth with respect to the untreated control mice (Figure 1 B and D). We continued the treatment beyond day 16 and monitored tumour growth in both groups of animals. At day 30, the size and weight of the tumours harvested from Defender®-treated mice were significantly reduced when compared to the untreated control mice (Figure 1C). Additionally, we observed the appearance of an extensive area of dead cells in the Defender®-treated tumour tissue, comprising approximately 23.94% of tumour, which was notably higher than the 10.81% measured for the untreated control mice (Figure 1E). The application of terminal deoxynucleotidyl transferase dUTP nick end labelling (TUNEL) staining with beta-actin further supported this finding, with a higher level of apoptotic cells present in the Defender®-treated tumour sections compared to the untreated sections (Figure 1F). These results collectively show that Defender® containing QV0 has a potent anti-cancer activity on the suppression of tumour growth in a PM animal model.

**Figure 1.**
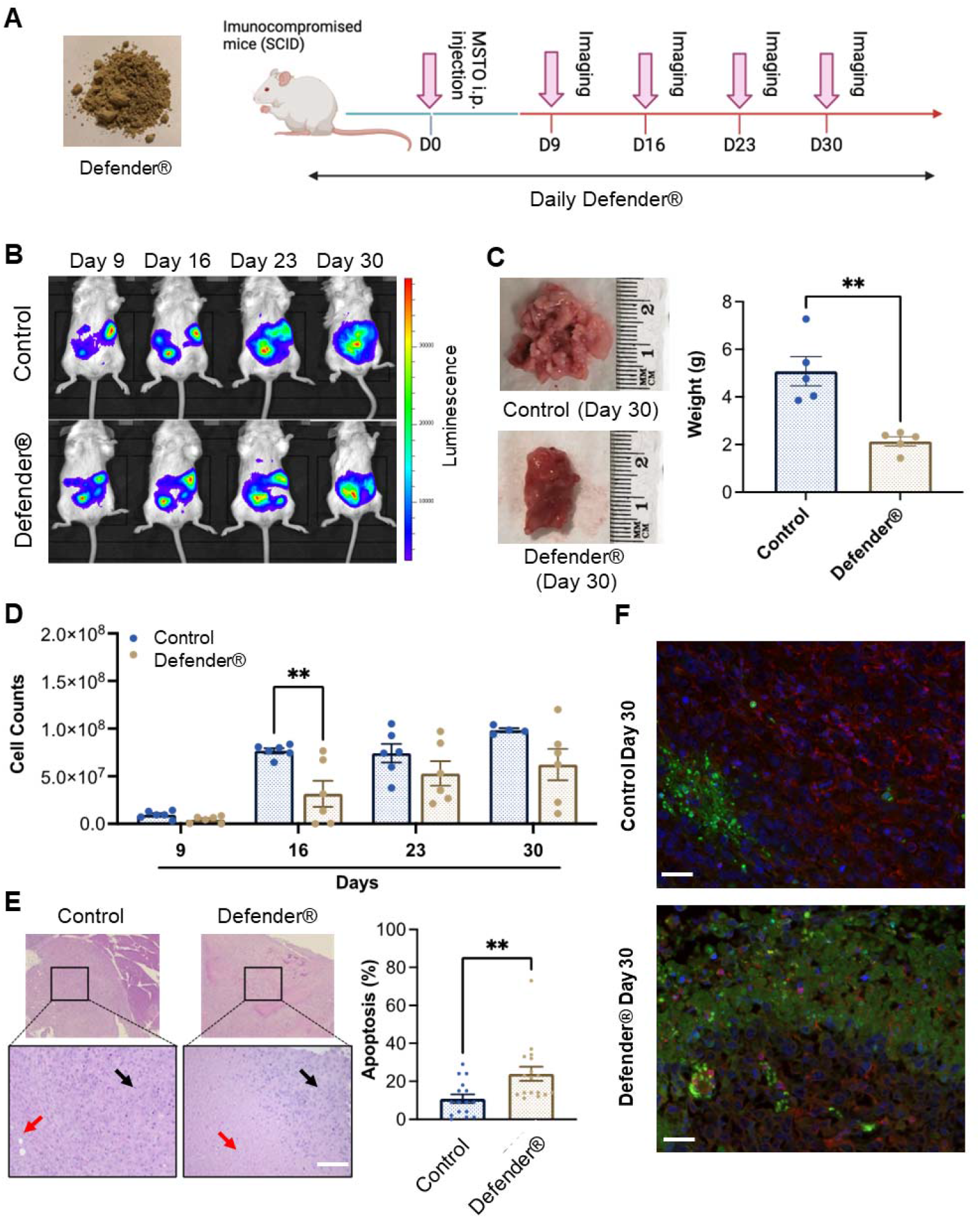
(A) SCID mice were inoculated with MSTO mesothelioma cells via intraperitoneal (i.p.) injection, followed by subsequent treatment with Defender® or saline administrated orally for 30 days. Pink arrows indicate the IVIS imager analysis schedule. (B) Representative images of animals treated with Defender® showing suppression of tumour growth with respect to the untreated control. (C) Defender® treatment significantly reduced tumour size and weight at harvesting with respect to the untreated control. (D) Tumour growth was quantified by total cell counts, as measured by the IVIS imaging system, which showed a reduction of tumour growth in animals treated with Defender® with respect to the untreated control. (E) Representative images showing an extensive area of dead cells (red arrows) in the Defender®-treated tumour H&E sections compared to the untreated control. Black arrows indicate the area of live cancer cells. (F) TUNEL staining showed enhanced apoptosis in Defender®-treated tumours. Green, red and blue staining corresponds to apoptotic cells (TUNNEL mix), live cells (Beta-actin) and nuclear DNA (DAPI), respectively. N=10 per group, **P<0.01

We also observed a significant increase in survival rate in tumour-bearing mice treated with Defender®, which was 7 days longer than the untreated control mice (Figure 2A). Additionally, we found that Defender®-treated mice had a reduced adverse effect index compared to the untreated control mice, which is measured based on criteria including a reduction in body weight, food intake and mobility, and development of bleeding or diarrhea (Figure 2B). More importantly, the Defender® administration did not induce the long-term systemic adverse effect in mice. For instance, the histological assessment of the stomach (gastric mucosa) showed no tumour involvement and no inflammatory features in both control and Defender®-treated animals. The spleens of control and Defender®-treated animals showed diffuse involvement by tumour dispersed as single cells. This resulted in a degree of disruption of white and red pulp, but architecture was preserved and discernible (Figure 2C). Tumour infiltration was observed in mice liver as small solid tumour nodules either with (20%) or without (66.7%) the treatment of Defender®, however the architectural distortion, cholestasis, ballooning, or fatty change which indicates the liver toxicity, was not detected (Figure 2D). Serum levels of alanine aminotransferase (ALT), aspartate aminotransferase (AST) and glucose were not changed post 30 days of the Defender® treatment (Figure 2E-G), which collectively demonstrated that the oral administration of Defender® for 30 days is did not induce liver toxicity in the PM animal model.

**Figure 2.**
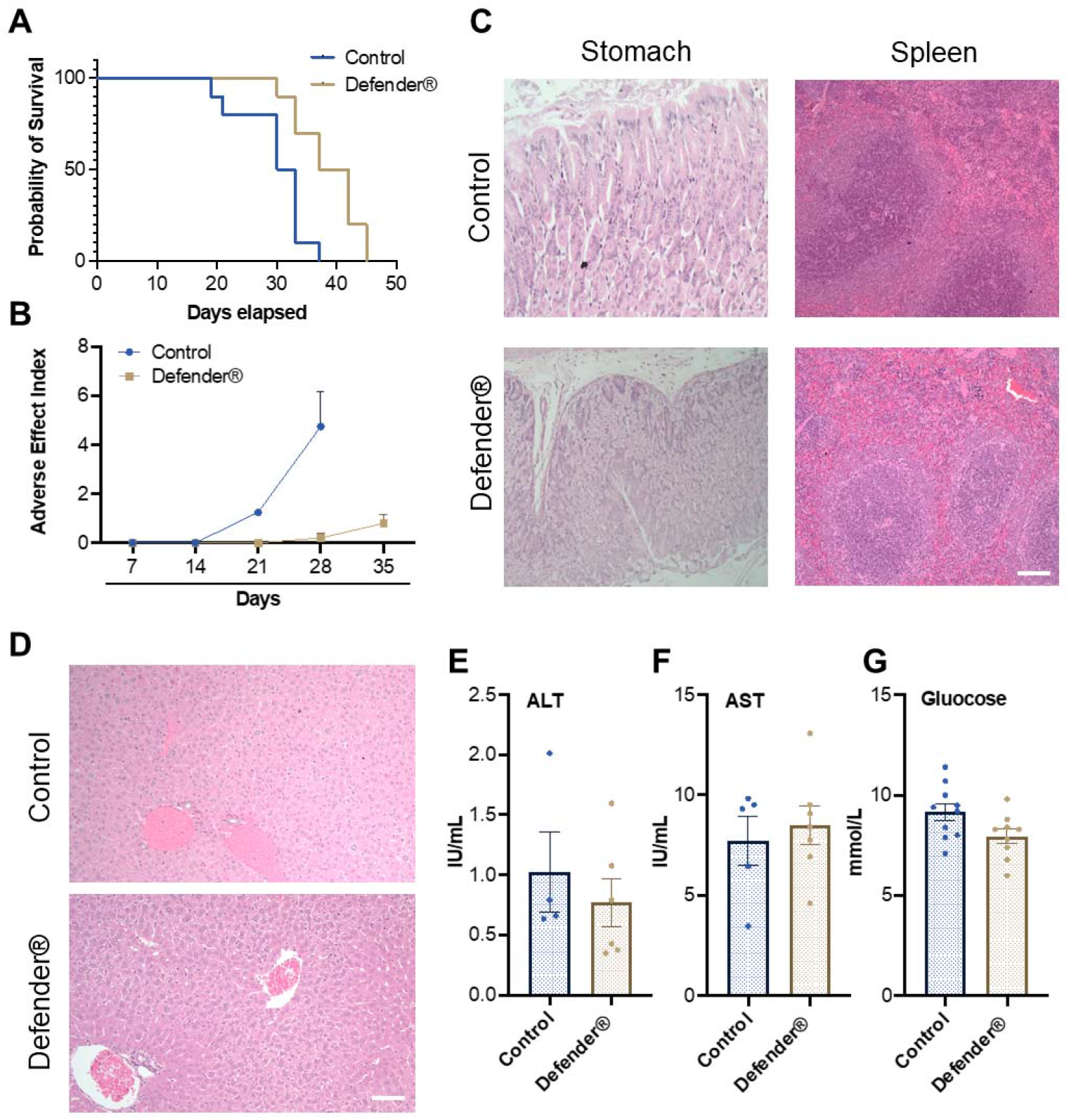
(A) Defender® treatment demonstrated an increased survival rate with an extended median survival. (B) Defender® treatment improved the adverse effect index after tumour bearing. (C) H&E staining showed that there were no major histological changes in stomach and spleen sections after Defender® treatment. (D) There was no architectural distortion, cholestasis, ballooning or fatty change in the liver of Defender®-treated animals. (E-G) No differences in serum ALT, AST or glucose concentration were observed post-Defender® treatment for 30 days. N=10 per group.

The food formula of QV0 defender® showed anti-cancer property in mesothelioma, we designed functional assays to understand the anti-cancer effects and mechanism of QV0, the Alamarblue® cell proliferation assay was used to assess the anti-proliferative effect of QV0 on 12 cancer cell lines; including eight mesothelioma, one breast cancer, one lung cancer, one gastric cancer, one colon cancer, and one non-cancer (MeT5A) cell(s). Results indicated that QV0 suppressed the growth of all 12 of the tested cancer cell lines in a dose-dependent manner at a concentration as low as approx. 0.02mg/ml (Figure 3A, Table 1). Interestingly, the non-cancer MeT5A cell line was substantially more resistant to QV0 treatment (0.12±0.01 g/mL) than any of the cancer cell lines (mean IC_50_ was 0.01±0.005g/mL; next highest IC_50_ was 0.02±0.002 g/mL) (Table 1). Furthermore, we found that the QV0-treated cancer cells at IC_50_ showed a significantly shorter migration distance in 24 hours (Welch two sample t-test, p=0.003, Figure 3B, C) and supressed the colony formation of cancer cells in 14 days (Figure 3D) (representative data of cells H2452 and MM05 are shown). However, in comparison to the normal cells, the treatment of QV0 for 48 hours did not induce any alterations to the cell cycle phases in the cancer cells (Figure 3E). Collectively, this finding suggests QV0 has the ability to inhibits cancer cell proliferation, cell migration and colony formation.

**Table 1.** IC_50_ values (concentration at which 50% of cells are viable) for each cell line treated with QV0.

| Cell line | $IC_{50}$ (g/mL) |
| --- | --- |
| MeT5A | $0.1234 \pm 0.0065$ |
| H28 | $0.0170 \pm 0.0017$ |
| H226 | $0.0100 \pm 0.0001$ |
| H2452 | $0.0162 \pm 0.0001$ |
| MSTO | $0.0173 \pm 0.0006$ |
| LnCap | $0.0077 \pm 0.0005$ |
| MKN45 | $0.0112 \pm 0.0011$ |
| MCF7 | $0.0087 \pm 0.0015$ |
| REN | $0.0079 \pm 0.00002$ |
| MM05 | $0.0184 \pm 0.0002$ |
| V40 | $0.0208 \pm 0.0019$ |
| H3122 | $0.0161 \pm 0.0009$ |
| AC29 | $0.0082 \pm 0.0004$ |

**Figure 3.**
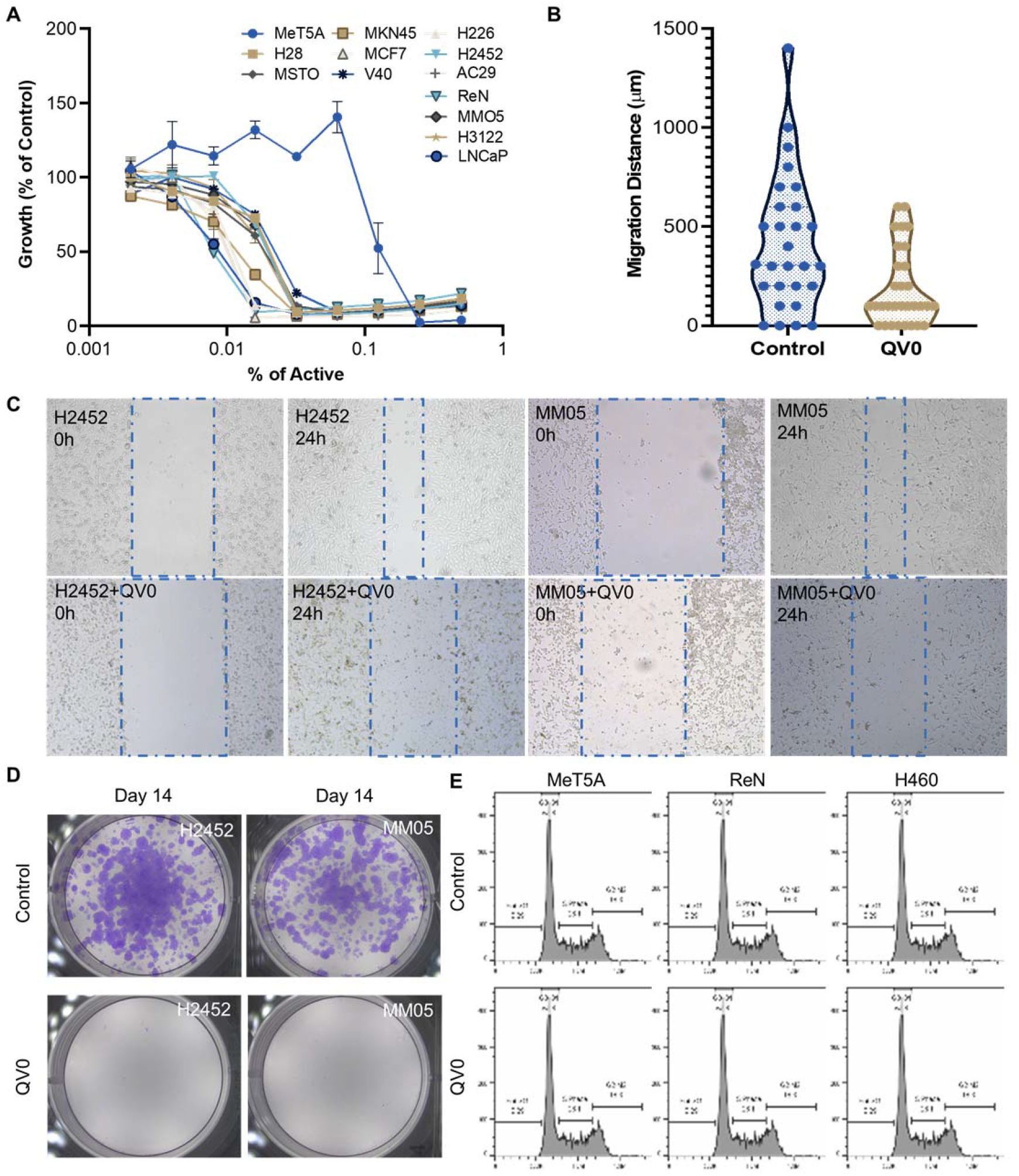
(A) QV0 suppressed growth of all tested cancer cell lines but not the non-cancer cell, MeT5A. The fluorescence intensity of the QV0-treated cells is presented as a percentage of the intensity of the untreated control cells. (B) QV0 significantly inhibited the tumour cell migration distance compared to the untreated control cells. (C) Representative images showing an inhibition of cancer cell (H2452, MM05) migration following QV0 treatment. (D) Representative images showing an evident suppression of colony formation in cancer cell lines, H2452 and MM05, following 14 days of QV0 treatment with respect to the untreated control cells. (E) Representative images depicting unaltered cell cycle profiles of normal mesothelial (MeT5A), mesothelioma cancer (Ren) and lung cancer (H460) cells at 48 hrs post-QV0 treatment. For each sample, 10000 events of single cells were counted and the cell cycle phases were subsequently analysed using FlowJo software. N=3 per each cell line.

We studied the effects of QV0 on cellular respiration and energy production in both non-cancer and cancer cells. The mitochondrial respiratory profile of the non-cancer cell line, Met5A, and mesothelioma cancer cell line, MSTO, was analysed using the Seahorse XF24 system following treatment with QV0. Live cells were sequentially injected with different mitochondrial respiration modulators, include oligomycin, phenylhydrazone (FCCP), Rotenone, and Antimycin (Figure 4A). The basal respiration measures the energetic demand of cells under basal conditions (Figure 4B) and maximal respiration represents the maximum capacity that the electro respiratory chain can achieve following the injection of FCCP (Figure 4C). ATP-linked respiration is reflected by the decrease in OCR following the injection of the ATP synthase inhibitor, oligomycin, which is the portion of basal respiration (Figure 4D). The remaining basal respiration not coupled to ATP synthesis after oligomycin injection represents the proton leak (Figure 4E)^18^.

**Figure 4.**
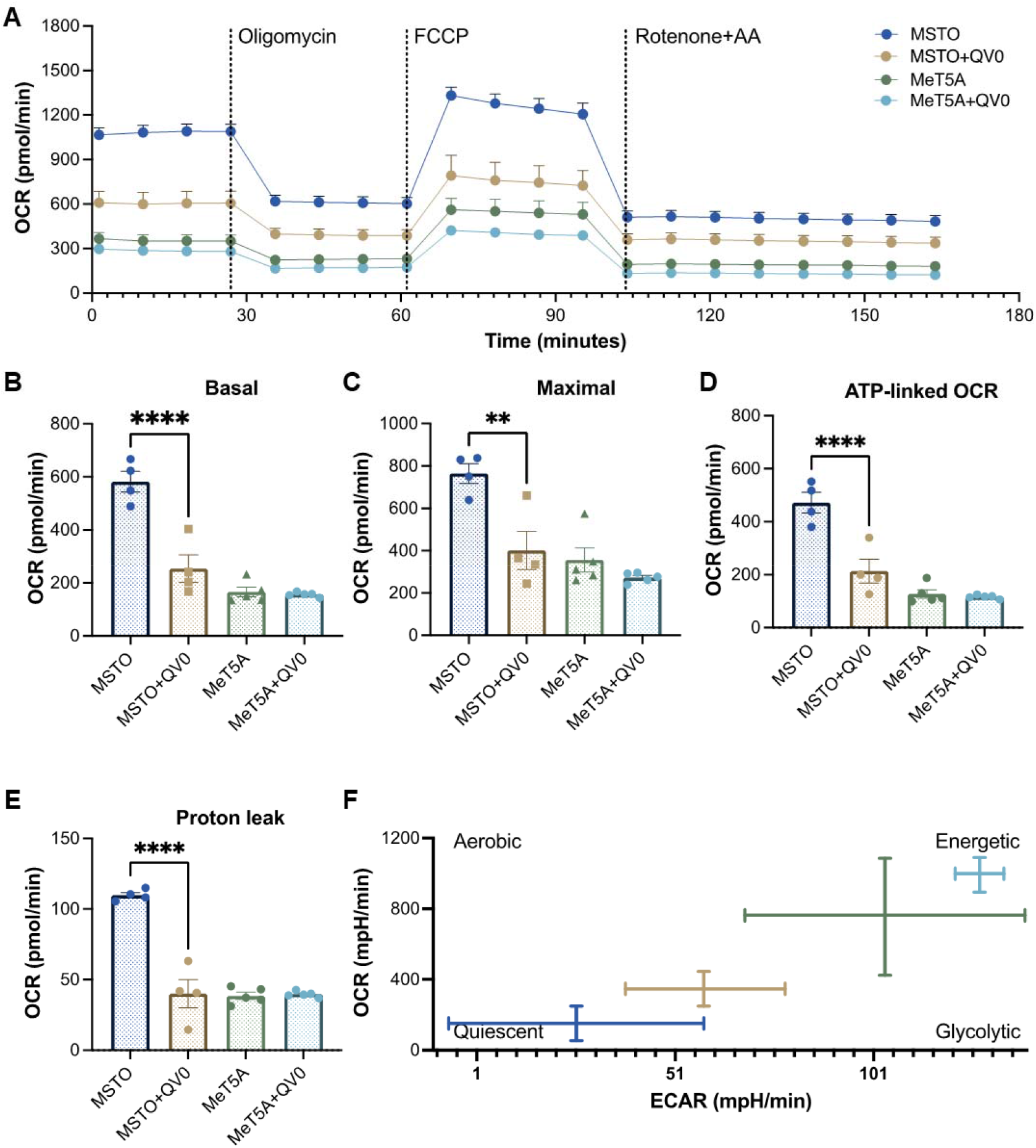
QV0 decreased PM cell energy demand and mitochondrial activity. (A) A representative profile of the mitochondrial stress test. Dotted vertical lines indicate the addition of 1 μM oligomycin, 0.5 μM FCCP, and 0.5μM Rotenone and Antimycin A. Graphs depicting (B) basal mitochondrial OCR, (C) maximal mitochondrial OCR, (D) ATP-linked OCR, (E) Proton Leak and (F) mitochondrial respiratory energy map (Ratio of OCR to ECAR). N=5 per group, **** P<0.0001

Our results indicate that the MSTO cells exhibited a higher level of mitochondrial activity with significantly higher basal, maximal and ATP linked respiration when compared to that of the non-cancer control, MeT5A (Figure 4A). Importantly, the treatment of QV0 significantly repressed the mitochondrial activity in MSTO cells, including basal (Figure 4B), maximal (Figure 4C), and ATP linked OCR (Figure 4D) as early as 24 hours post the QV0 treatment. Moreover, the proton leak, which can be a sign of mitochondrial damage, was significant reduced in QV0-treated MSTO cells at 24 and 48 hours, suggesting an impaired mitochondrional function (Figure 4E, S2). However, it was of interest that QV0 exerted minimal effect on mitochondrial function in the non-cancer cell, MeT5A (Figure B-E). The basal extracellular acidification rate (ECAR) was plotted against OCR in Figure 4F, where the energetic (MSTO) and quiescent (MeT5A) bioenergetic profiles were demonstrated. A shift in bioenergetics was observed for MSTO following QV0 treatment after 24 hours, with cells becoming less energetic and more quiescent as the ATP production pathways were inhibited.

We then examined the chemical and antioxidant properties including total phenolic content and ferric antioxidant power (FRAP) of QV0, Defender® and animal plasma samples after oral Defender® treatment. The result demonstrated that Defender® is a rich source of phenolic compounds, which are higher than those of QV0, and plasma has the lowest level of phenolic compounds. Noticeably, ferric antioxidant power of QV0 is significantly higher than that of Defender® and plasma (Table 2). Quercetin and Kaempferol are two phenolic compounds identified in QV0 and QV0 has significantly higher levels of these compounds as compared to that of the Defender®. Of note, Quercetin and Kaempferol have not been identified in the plasma, however, there are two compounds (peaks 1 and 2, Figure S1) have been identified in QV0, Defender® and plasma. QV0 has the highest levels of the two unknown compounds, followed by Defender® and plasma. In addition, the scanning results (Figure S1) revealed that there are over 30 major peaks can be observed in both QV0 and Defender®, meaning there are over 30 major individual phytochemicals and most of these compounds have not been identified.

**Table 2.**
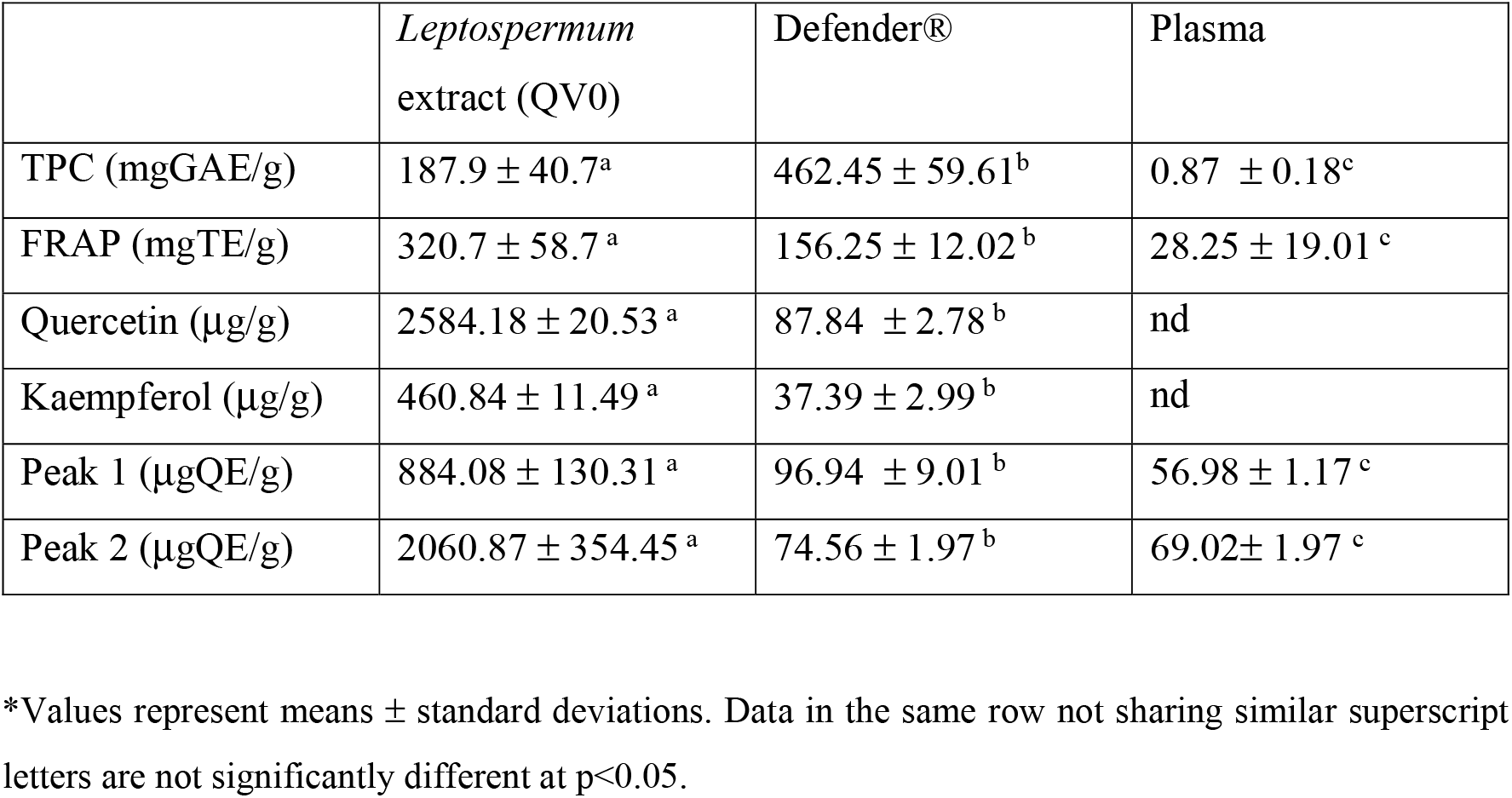
Total phenolic content, antioxidant activity, and some phytochemicals in *Leptospermum* extract (QV0), Defender®, and plasma samples.

|  | *Leptospermum*<br>extract (QV0) | Defender® | Plasma |
| --- | --- | --- | --- |
| TPC (mgGAE/g) | 187.9 ± 40.7<sup>a</sup> | 462.45 ± 59.61<sup>b</sup> | 0.87 ± 0.18<sup>c</sup> |
| FRAP (mgTE/g) | 320.7 ± 58.7<sup>a</sup> | 156.25 ± 12.02<sup>b</sup> | 28.25 ± 19.01<sup>c</sup> |
| Quercetin (µg/g) | 2584.18 ± 20.53<sup>a</sup> | 87.84 ± 2.78<sup>b</sup> | nd |
| Kaempferol (µg/g) | 460.84 ± 11.49<sup>a</sup> | 37.39 ± 2.99<sup>b</sup> | nd |
| Peak 1 (µgQE/g) | 884.08 ± 130.31<sup>a</sup> | 96.94 ± 9.01<sup>b</sup> | 56.98 ± 1.17<sup>c</sup> |
| Peak 2 (µgQE/g) | 2060.87 ± 354.45<sup>a</sup> | 74.56 ± 1.97<sup>b</sup> | 69.02 ± 1.97<sup>c</sup> |

\*Values represent means ± standard deviations. Data in the same row not sharing similar superscript letters are not significantly different at p<0.05.

## Discussion

Chemotherapy is one of the most commonly administered treatments for mesothelioma, however there are many adverse side effects associated with its use on PM patients, such as hair loss, nausea, vomiting, diarrhea, fatigue, and immune suppression^19^. The current combination of ipilimumab and nivolumab immunotherapy has similar response rate to chemotherapy but risk of immune mediated adverse events is modest. The discovery of improved treatment options with minimal side effects is urgently needed. Natural extracts have continuously proven to be an important and rich source of anti-cancer therapies^20^, however limited studies have investigated their potential utility for the treatment of mesothelioma. In the present study, we investigated for the first time the anti-cancer effect of *Leptospermum* extract (QV0) in PM and found that the food formula of QV0 (Defender®) can suppress mesothelioma growth in a mouse model. Defender®-treated animals showed a reduction in tumour volume and improved health index, which was associated with an extended life expectancy of up to 7 days when compared to untreated animals. During the 30-day treatment period involving oral administration of Defender®, the animals showed no evident signs of liver toxicity, nor was there an increase in their blood glucose level. Additionally, the histology of the spleen, liver and stomach post-Defender® treatment was assessed by a pathologist, whereby no adverse side effects were observed. This finding does not reflect that of manufactured medicines, such as chemotherapy drugs, which typically induce multiple adverse side effects. Plant extracts such as those used in Traditional Chinese Medicine (TCM) have been practiced and developed over thousands of years^21^, however some have been known to cause liver toxicity^22^. Given that our study showed no evident signs of liver toxicity following Defender® treatment, this suggests that the potential use of Defender® for the treatment of PM would be a safer alternative to conventional TCM and provides rationale for further testing of Defender® in prospective human clinical trial studies.

In addition to the animal study, we investigated the anti-cancer activity of QV0 in PM cells. Our result indicated that QV0 has the ability to suppress cell proliferation and migration in mesothelioma cells (H28, MSTO, VMC40, H226, H2452, REN, MMO5 and AC29), breast cancer (MCF7), prostate cancer (LnCAP) and lung cancer (H3122) cell lines. The IC_50_ indicated that an immobilised mesothelial cell, MeT5a, was less sensitive to the QV0 treatment at the effective concentration (0.2g per 100 ml), whereas all cancer cells were found to be sensitive at QV0 concentrations ranging from 0.01 to 0.02g per 100ml. We concluded that QV0 has no significant toxicity in non-cancer cells. A clonogenic assay indicated that QV0 significantly suppressed mesothelioma cell colony formation. In a mesothelioma preclinical model, we also established that Defender®-treated animals showed significant tumour suppression. Although QV0 is derived from the *Leptospermum* plant to the best of our knowledge, there are no scientific reports based on the anti-cancer properties of manuka honey-based products for mesothelioma. Therefore, our study is the first of its kind to demonstrate that a plant extract (QV0) derived from the *Leptospermum* tree exhibits anti-cancer properties in both a mesothelioma animal model and mesothelioma cell lines. Our result indicated that in Defender®-treated animals, the harvested tumours appeared to have a significant increase in cell death when compared to the untreated control animals. The tunnel assay confirmed that QV0 treatment induced mesothelioma cell apoptosis. This finding is concordant with similar studies by Navanesan et al. who demonstrated the leptospermum sub spp. (similar species of QV0) Javanicum and flavescens are capable of inducing cell apoptosis and supressing the metastatic potential of human lung carcinoma cells^23,24^.

The mechanisms of apoptosis are highly complex but commonly caused by two main pathways; the extrinsic death receptor pathway and intrinsic mitochondrial pathway. In the present study, QV0 treatment did not trigger cell-cycle arrest, suggesting apoptosis is not caused by cell cycle profile changes. We then performed the mitochondrial stress test to measure the mitochondrial function, specifically mitochondrial OCR. The results demonstrated a significant reduction of the basal, maximal and ATP-linked OCR in the QV0-treated cancer cells, with an evident shift in metabolic potential from energetic to quiescent as early as 24 hours post-treatment. The suppression of proton leaks also indicated the mitochondrial damage after QV0 treatment. These findings suggest that QV0 inhibits the mitochondrial OCR in mesothelioma cells, causing mitochondrial dysfunction-induced apoptosis. Our findings are in agreement with those of a study by Amran et al., which demonstrated that Tualang honey inhibits cell proliferation and induces cell apoptosis with reduced mitochondrial membrane potential in the human breast cancer cell lines, MCF-7 and MDA-MB-231^25^. Moreover, we found that the anti-cancer effects of QV0 on proliferation, apoptosis and mitochondrial function were cancer cell-specific, with QV0 having no effect on the non-malignant cells, thus suggesting its potential to be used as a novel anti-cancer drug that induces minimal damage or alteration to healthy non-malignant cells. Overall, these comprehensive studies conclude that as a natural plant extract, QV0 possesses desirable anti-cancer properties, both *in vitro* and *in vivo*, which is associated with and/or mediated by mitochondrial dysfunction-related apoptosis.

Anti-cancer properties of QV0 and Defender® can be attributed to their high level of polyphenols, which possess strong antioxidant activity. Levels of polyphenols in QV0 and Defender® are higher than that in the ginseng root extract^26^ and selected Chinese and Mexican medicinal plant extracts ^27,28^. Although QV0 contains less than 50% of polyphenols compared to that of the Defender®, the antioxidant activity of QV0 is significantly higher (double) than that of Defender®, revealing that phenolic compounds in QV0 exhibit potent antioxidant activity compared to those of the Defender®, which only contains 5% of QV0. Of note, there are over 30 major individual compounds observed from Figure S1, however there are only two compounds identified: Quercetin and Kaempferol. These compounds are known to induce cytotoxic effects on cancer cells through several mechanisms, such as apoptosis, cell cycle arrest at the G2/M phase, and downregulation of epithelial-mesenchymal transition (EMT) – related markers ^29,30^. Over 28 individual compounds are yet to be identified and tested for their anti-cancer properties. Therefore, future studies are warranted to isolate and characterise these compounds, and to subsequently investigate their potential anti-cancer properties. Of note, level of antioxidant phenolic compounds is low in the plasma of the treated animals for over the 30 days, revealing that most phenolic compounds are absorbed through the whole process of metabolism into the bloodstream. In addition, there are two peaks (peaks 1 and 2, Figure S1) were observed in the QV0, Defender and plasma, these peaks would be likely the key compounds that involved in anti-cancer properties. Future studies are recommended to identify these compounds and their mechanisms on anti-cancer properties.

## Summary

PM is an aggressive malignancy of the lung lining with limited effective treatment options. In the present study, we have shown the promising anti-cancer potential of the tree *Leptospermum polygalifolium*-derived natural product, QV0 and Defender®. Specifically, this study demonstrates that QV0 exerts an inhibitory effect on PM tumour cell growth and improves host survival in a PM mouse model. These exciting findings provide an essential foundation and rationale for early-stage clinical trials, and we believe that prospective translational research will facilitate the successful implementation of QV0 in the clinical setting as a novel treatment option that will ultimately benefit PM patients.

## Materials

*Leptospermum polygalifolium* extract (QV0, P116949.AU) and dietary supplement (Defender®) were supplied by the Quality Global Supply Australia Pty Ltd, Tuggerah, NSW, which produces natural supplements and honey-based products. QV0 was prepared from *Leptospermum polygalifolium leaves* and small stems using aqueous extraction, following by spray drying to obtain powdered extract. For feeding animals, QV0 was prepared in the form of a dietary supplement (Defender®), which includes 5% of QV0, 15% of citrus pomace powder and 80% of honey.

## Methods

### In vitro

#### Cell proliferation assay

Briefly, 2,500 cells were seeded in 96-well culture plates in 200 μL medium per well overnight. Cells were treated with QV0 (IC_50_) for 72 hours, followed by the subsequent addition of 20 μL Alamarblue® (50 mL PBS containing 0.075 g Resazurin, 0.0125 g Methylene Blue, 0.1655 g Potassium hexacyanoferrate (III), 0.211 g Potassium hexacyanoferrate (II) trihydrate, filter-sterilised, and stored at 4°C in the dark). The cells were then incubated for 4 hours at 37°C as described ^31^. Fluorescence intensity was measured at 590 nm with 544 nm excitation using a FLUOstar Optima (BMG LabTech, Ortenberg, Germany). Fluorescence intensity was calculated as a percentage of the total intensity of the untreated control cells. Experiments were performed 3 times with 3 replicates each time, except for experiments involving slow-growing non-cancer primary fibroblasts that were performed 4 times with 2 or 3 replicates.

#### Cell migration assay

Cell migration of various cell lines were measured using a scratch (wound-healing) assay. Briefly, cells were plated in 24-well plates and at 24 hours post-seeding, 10 μg/mL camptothecin (Sigma-Aldrich) was added to stop cell proliferation; at the same time, a cross-shaped scratch was made using a 200 μl plastic pipette tip. At 12 and 24 hours post-scratch, microscopic imaging was carried out with a 20X objective (Leica DMi1). Each experiment group was performed in duplicate.

#### Clonogenic assay

Cells of each cell line were seeded in 6-well culture plates at a seeding density of 2,500 cells/well. QV0 (IC_50_) was added at 2 hours post-seeding and culture plates were incubated for 10-14 days at 37°C. Cells were then fixed with 70% ethanol and stained with 0.1% crystal violet before being photographed for colony counting using a ZEISS Stemi508 microscope.

#### Cell Cycle analysis

Mesothelioma cells were treated with QV0 (IC_50_) and at 48 hours post-treatment, the cells were harvested and washed 3 times with phosphate-buffered saline (PBS). The cells were subsequently fixed with 70% ethanol for at least 30 mins. For cell cycle analysis, fixing solution was removed and cells were treated with 0.01 % RNase (10 mg/ml, Sigma-Aldrich), 0.05 % propidium iodide (PI)(Sigma-Aldrich) in PBS for 30 min at 37°C in the dark. The cell cycle distribution was determined on a CytoFLEX flow cytometer (Beckman Coulter, Miami Lakes, FL) within 30 minutes. The flow cytometer was calibrated using calibration beads according to the manufacturer’s instructions (CytoFLEX, Beckman). The flow cytometer was routinely operated at the slow flow rate setting (μL sample/minute), and the data acquisition for a single sample typically took 3-5 mins. For each sample, 10,000 events of single cells were counted and the cell cycle was analysed using FlowJo software (Ashland, OR, USA).

#### Seahorse extracellular flux analysis

The seahorse XF24 Extracellular Flux Analyzer (Agilent, CA, USA) was used to measure the respiration activity of mesothelioma cell. Cells were seeded at a seeding density of 8 × 10^4^ cells per well in an XF24 plate overnight and treated with and without QV0 for 24 hours. The mitochondrial stress test was performed according to the manufacturer’s instructions. Briefly, 1 μM oligomycin (oligo), 0.3 μM FCCP, and 1 μM Rotenone and Antimycin A were added and the relative levels of basal, maximal respiration and reserved mitochondrial capacity were calculated based on OCR data obtained from the Mito stress tests using Seahorse Wave software for XF analyzers (Agilent, CA, USA).

### In vivo

#### PM Xenograft mouse model

To study the *in vivo* response of QV0, a food formula ‘Defender®’ (consisting of honey as a base and containing QV0) was used as supplement to feed the animals. A total of twenty SCID mice (8 week old females) were injected with 1 × 10^6^ human mesothelioma cells (MSTO-211H) pre-transfected with a stable pGL4-51lu luciferase construct for visualisation of tumour grown. Animals were then evenly separated into two groups (control and Defender®). In the Defender® group, animals were treated 5mg/mouse/20g body weight with Defender® as a daily supplement, which was administrated orally using an oral gavage. The tumour was measured on a weekly basis by luciferin for visualisation (150mg/kg). Images were taking for a period of at least 30 days by IVIS (PerkinElmer, Waltham, USA).

#### Histological assessment

Animal tumour, liver, spleen and stomach tissues were embedded in paraffin. Multiple 4μm sections were stained with haematoxylin and eosin (H&E) (Sigma-Aldrich) for general histological analysis by two pathologists.

#### TUNEL assay

An *In situ* cell death detection kit (Roche, Basel, Switzerland) was used for the detection and quantification of tumour cell apoptosis. Briefly, formalin fixed tumour tissue sections were dewaxed according to standard procedures and then incubated in 0.1M citrate buffer PH 6.0 at 70 °C for 1 hour. The slides were blocked with Tris-HCL, 0.1M PH 7.5, containing 3% BSA and 20% normal bovine serum for 30 mins at room temperature, followed by the addition of 50 uL of TUNEL reaction mixture to the slides and subsequent incubation for 60 mins at 37°C in a humidified atmosphere in the dark.

#### Liver toxicity test

Liver toxicity was assessed by aspartate aminotransferase (AST) and alanine aminotransferase (ALT) serum concentrations using commercial assays (MAK055 and MAK052, Sigma-Aldrich). All experimental work was performed as per kit instructions.

#### Analysis of total phenolic content (TPC)

The level of TPC was determined by the Folin-Ciocalteu method (AOCS, 1990), modified for the microplate. Water was used as the blank and gallic acid was used as the standard for a calibration curve. 15 μL of sample, standard or blank was placed into 24-well microplates, followed by the addition of 240 μL of diluted Folin-Ciocalteu (FC) phenol reagent (6.25 %). The mixture was incubated in the dark for 10 mins at room temperature, followed by the addition of 15 μL of 20% sodium carbonate. The mixture was then incubated for a further 20 mins in the dark, and absorbance was measured at 765 nm using a FLUOstar Optima microplate reader (BMG LabTech, Ortenberg, Germany). The TPC was calculated based on the calibration curve and expressed as milligrams of gallic acid equivalent per gram of sample (mgGAE/g).

#### Ferric Reducing Antioxidant Power (FRAP)

Antioxidant activity of the sample was evaluated using the ferric reducing antioxidant power assay (FRAP) according to the previously published method [1] with modification for the microplate. Briefly, a FRAP working solution was prepared freshly by mixing three reagents: (A) 300 mM acetate buffer (PH 3.6), (B) 10 mM TPTZ (2,4,6 tripyridyl-s-triazine) in 40 mM HCL solution and (C) 20mM Ferric Chloride. Trolox was used as the standard for a calibration curve. 50 uL of the sample, blank, or standard was placed into 24 well microplates, followed by the addition of 300 uL of FRAP working solution. The mixture was incubated for 30 min at room temperature and absorbance was measured at 593 nm using a FLUOstar Optima microplate reader (BMG LabTech, Ortenberg, Germany). Antioxidant activity of plasma was expressed as milligrams of Trolox equivalent per gram of extract (mgTE/g).

#### Preparation of Defender® solution and plasma for analysis

QV0 powder (5g) was diluted in water (100mL), then filtered through a 0.45 μm nylon membrane prior to analysis. Animal plasma protein was precipitated by adding 12% trichloroacetic acid (TCA) at the ratio of animal plasma sample to TCA of 1:2 (v/v) and was mixed by vortexing. The mixture was then centrifuged at 2500 rpm for 15 min. The supernatant was used for further analysis.

#### Scanning for major phytochemicals

Major phytochemicals in *Leptospermum* extract and plasma were determined using a Shimadzu HPLC system (Shimadzu, Japan) fitted with a reverse phase column (Luna 5u Phenyl-Hexyl 250 x 3.00 mm 5u micron) (Phenomenex) maintained at 35 °C by a column oven (CTO-20A, Shimadzu) with photodiode array detector (SPD-M40). The mobile phase consisted of 0.1% formic acid (Solvent A) and absolute acetonitrile (Solvent B). An auto injector (SIL-20A) was used to inject 25 μL sample volumes onto the HPLC at a flow rate of 0.7 mL/min with a gradient elution schedule as follows: 0-10 min, 0% B; 10-45 min, 40% B; 45-60 min, 60% B; 60-70 min, 60% B; 70-80 min, 0% B and 80-85 min, 0% B. Kaempferol was used as the standard for quantification of Kaempferol. Quercetin was used as the standard for a calibration curve for quantification of Quercetin and two unknown phytochemicals and the results were expressed as milligram of quercetin per gram of sample (μgQE/g).

#### Statistical analysis

For the proliferation assays, the QV0 IC50 concentration at which 50% of cells were viable was calculated by modelling cell response to QV0 treatment using a sigmoid function^32^ as described previously^31^. Briefly, the sigmoid function used to predict cell proliferation, *y*, was:

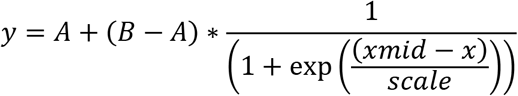

where *A* is the left asymptote (cell response at QV0 treatment concentration of 0), *B* is the right asymptote (cell response at highest QV0 treatment concentration), *xmid* is the transition point (IC50) of the cells treated with QV0, *scale* is an x-axis scale parameter impacting slope of the transition, and *x* is log10 of the QV0 treatment concentration (thus rendering the curve symmetrical and suitable for modelling using log-likelihood). The best fitting parameters for a given model were determined by the maximum log likelihood method, using the optimx package^33^in R^34^. IC50 (concentration at which 50% of cells are viable) was calculated as the sigmoidal transition point resulting from the model having the best fitting parameters. IC50 standard deviation was calculated as the standard deviation of the transition points for each experiment modelled individually as a sigmoid function.

The cell cycle profile of a cell line after QV0 treatment was compared to that of the same cell line having no QV0 treatment, in the following way. The cell cycle profile is the percentage of cells in each cell cycle phase. The differences between the QV0-treated and non-treated cells were calculated. Student’s t-test in R^34^ was used to determine whether the differences were zero. ANOVA and paired t-test were also used in this study with significance set at P<0.05.

## Authors contribution

HS and YC conceived the project, conduct the experiments, and prepared the manuscript, Le Z, T.K Y, Ling Z, Thuy P and QV assist in experiments, ER and HS performed the data analysis, KL and Sonja K prepared sections and performed histology assessment, Le Z prepared figures, Le Z, HK, BJ and Steven K edited manuscript. All authors contributed to the manuscript and approved the submitted version.

## Acknowledgement

This project is partially supported by Quality Global Supply Pty Ltd

---

**Figure S1.**
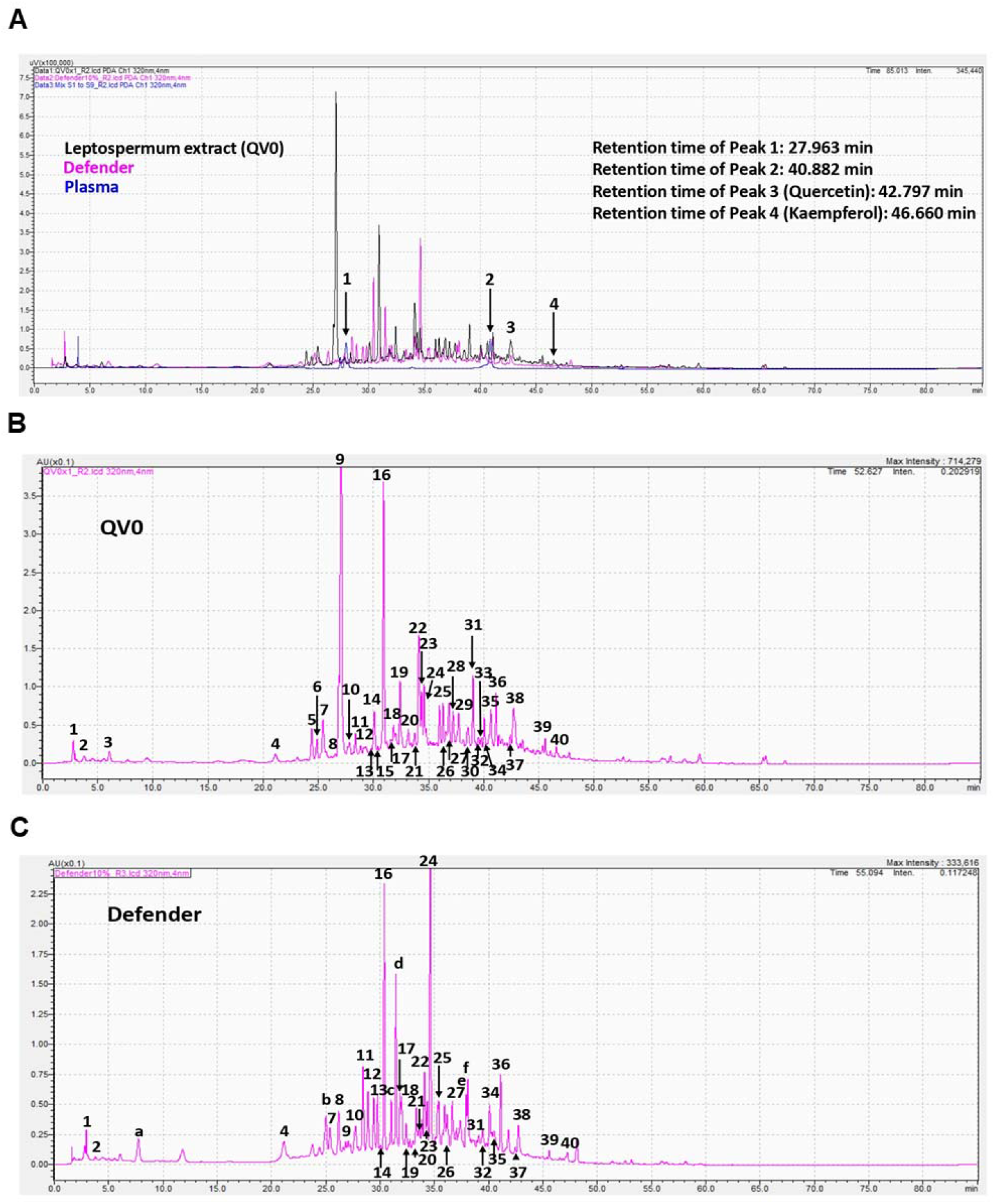
Chromarograms of Leptospermum extract (QV0), Defender, and Plasma (A); QV0 (B); and Defender (C) measured at 320nm using a PDA detector.

